# Heavy chain-1 of inter-α-inhibitor has an integrin-like structure with immune regulatory activities

**DOI:** 10.1101/695700

**Authors:** David C. Briggs, Alexander W.W. Langford-Smith, Thomas A. Jowitt, Cay M. Kielty, Jan J. Enghild, Clair Baldock, Caroline M. Milner, Anthony J. Day

## Abstract

Inter-α-inhibitor (IαI) is a proteoglycan essential for mammalian reproduction that also plays a less well-characterised role in inflammation. IαI is composed of 2 homologous ‘heavy chains’ (HC1 and HC2) covalently attached to chondroitin sulphate on the bikunin core protein. Prior to ovulation HCs are transferred onto the polysaccharide hyaluronan (HA), thereby stabilising a matrix that is required for fertilisation. Here we show that human HC1 has a structure similar to integrin β-chains and contains a functional MIDAS (metal ion-dependent adhesion site) motif that can mediate self-association of heavy chains, providing a mechanism for matrix crosslinking. Surprisingly, its interaction with RGD-containing integrin ligands, such as vitronectin and the latency-associated peptides of TGFβ, occurs in a MIDAS/cation-independent manner. However, HC1 utilises its MIDAS motif to bind to, and inhibit the cleavage of, complement C3, thus identifying it as a novel regulator of innate immunity through inhibition of the alternative pathway C3 convertase.

**Abbreviations:** ADPs, atomic displacement parameter; AUC, analytical ultracentrifugation; CMG2, capillary morphogenesis protein-2; COC, cumulus-oocyte complex; CS, chondroitin sulphate; FB, complement factor B; FnIII; fibronectin type III; HA, hyaluronan; HC, heavy chain; HC•HA, covalent complex of HC with HA; IαI, inter-α-inhibitor; ITGA, integrin α-chain; ITGB, integrin β-chain; LAP, latency associated peptide; LLC, large latent complex; LTBP, latent TGFβ binding protein; MIDAS, metal ion-dependent adhesion site; PαI, pre-α-inhibitor; PTX3, pentraxin-3; rHC1, recombinant HC1; SAXS, small-angle X-ray scattering; SHAP, serum-derived HA binding protein; SLC, small latent complex; TEM8, tumour endothelial marker-8; TGFβ, transforming factor β; TSG-6, tumour necrosis factor-stimulated gene-6; TSG-6•HC, covalent complex of TSG-6 and HC; vWFA domain, von Willebrand Factor A domain.

## Introduction

Inter-alpha-inhibitor (IαI) is an unusual plasma proteoglycan, comprised of 3 protein chains covalently linked via a chondroitin-sulphate (CS) glycosaminoglycan (GAG) chain (Enghild et al., 1989); see Supplementary Figure 1A. The CS chain is attached via a typical tetrasaccharide linkage to the core protein, bikunin. The protein products of the ITIH1 and ITIH2 genes, termed heavy chain 1 (HC1) and HC2, are covalently attached via ester bonds linking their C-termini to C6 hydroxyl-groups on the N-acetyl galactosamine sugars within the CS chain (Enghild et al., 1993; Morelle et al., 1994). HC2 is positioned closer to bikunin than HC1, and the two HCs are attached to sugars 1-2 disaccharides apart (Enghild et al., 1999; Ly et al., 2011).

HC1 and HC2 are approximately 80 kDa in size and share ∼39% sequence identity. They are synthesised with C-terminal pro-domains (of 239 and 244 amino acid residues, respectively) that are cleaved when the HCs are covalently attached to the bikunin CS chain (Kaczmarczyk et al., 2002; Zhuo et al., 2004); HC3 (ITIH3; 54% identical to HC1) can also link to the bikunin CS proteoglycan (Supplementary Figure 1A), to form pre-alpha-inhibitor (PαI) (Enghild et al., 1989, 1991), and there is evidence that the related HC5, and likely HC6 (but not HC4), can also become attached to CS in this way (Day and Milner, 2019; Martin et al., 2016). All HCs are predicted to contain a single von Willebrand factor type-A (vWFa) domain, which makes up roughly the central one-third of the amino acid sequence. The flanking sequences are not homologous to any known domain.

IαI plays a critical role in mammalian reproductive biology such that female mice with the bikunin gene deleted, and consequently lacking IαI and PαI, are infertile (Sato et al., 2001; Zhuo et al., 2001). This is due to the impaired formation of the cumulus extracellular matrix that normally drives the expansion of the cumulus-oocyte-complex (COC). This elastic matrix (Chen et al., 2016) protects the oocyte during the expulsion of the COC from the follicle and also provides a large surface area facilitating sperm capture *in vivo* (Nagyova, 2015; Russell and Salustri, 2006). The cumulus matrix is rich in the non-sulphated GAG hyaluronan (HA), where this high molecular weight polysaccharide becomes modified by the covalent attachment of HC1, HC2 and HC3 (Mukhopadhyay et al., 2001). Here, TSG-6, a protein that is expressed by the cumulus cells, plays a catalytic role in transferring the HCs from the CS chains of IαI and PαI onto HA to form HC•HA (aka SHAP-HA) complexes (Day and Milner, 2019; Rugg et al., 2005). TSG-6 also mediates the formation of HC•HAs during inflammation, when IαI/PαI leak into tissues from the circulation.

The covalent attachment of HCs changes the physical properties of HA. For example in synovial fluid from rheumatoid arthritis patients, where on average 3 to 5 HCs are attached to an HA chain of ∼2 MDa, the polysaccharide is more aggregated compared to unmodified HA (Yingsung et al., 2003); this has been attributed to crosslinking of HC•HA complexes via interactions between HCs based on their apparent associations visualised by electron microscopy. Given that HC1, HC2 and HC3 can all be transferred onto HA during arthritis (Zhao et al., 1995) such crosslinking could be mediated by homotypic and/or heterotypic HC-HC interactions. Irrespective of the mechanism, the formation of HC•HA in arthritic joints enhances the binding of HA to its major cell surface receptor, CD44, on leukocytes (Zhuo et al., 2006), but it is unknown whether this, or indeed the altered hydrodynamic properties of the modified HA (Baranova et al., 2014), are part of a protective process or contributing to pathology (Day and Milner, 2019).

In some contexts, HC•HAs can be crosslinked by the binding of HCs to the octameric protein pentraxin-3 (PTX3) (Baranova et al., 2014), where, for example, the multivalent nature of PTX3 is essential to the stabilisation of the cumulus matrix (Inforzato et al., 2008). Either deletion of PTX3, or loss/impairment of TSG-6’s HC transferase activity in mice, also leads to the failure of COC expansion and, hence, infertility (Briggs et al., 2015; Fülöp et al., 2003; Ochsner et al., 2003; Salustri et al., 2004). Here the cooperation between HA, IαI, PTX3 and TSG-6 (Baranova et al., 2014) leads to the formation of an elastic tissue, which is the softest described to date with a Young’s modulus on the order of 1Pa (Chen et al., 2016).

IαI has been implicated as a regulator of innate immunity having been shown to be an inhibitor of the complement system, affecting the alternative, classical and lectin activation pathways (Adair et al., 2009; Garantziotis et al., 2007; Okroj et al., 2012). The inhibition of the alternative and classical pathways of complement is thought to be dependent on the HCs rather than bikunin (Garantziotis et al., 2007; Okroj et al., 2012); i.e. both HC1 and HC2, isolated from native IαI, had inhibitory activities in haemolytic assays (Okroj et al., 2012), however, the mechanism has not been determined. In the case of the alternative pathway, IαI was found to inhibit the enzymatic cleavage of factor B (FB) to Bb, which occurs during the formation of the C3 convertase (C3bBb), and there is some evidence that IαI may interact with complement C3 (Garantziotis et al., 2007; Okroj et al., 2012).

IαI has also been found to bind to vitronectin (Adair et al., 2009), a multifunctional plasma and matrix protein that, as a well as being a regulator of complement system terminal pathway, also mediates binding to α_V_ integrins (Preissner and Reuning, 2011). Vitronectin’s integrin-binding activity has an important role in epithelial repair in the context of lung homeostasis and the adhesion and migration of epithelial cells was promoted by its interaction with IαI (Adair et al., 2009); moreover, IαI-deficient mice had impaired recovery in experimental lung injury. The association between IαI and vitronectin is reported to be of high affinity and inhibited by RGD peptides, implicating IαI’s vWFa domain in the interaction.

In order to explore and better explain the functions of HCs, we undertook structural and biophysical characterisation of the prototypical heavy chain, HC1. Here we present the crystal structure of HC1, and reveal that HC1 can form metal ion-dependent homodimers, which require a functional MIDAS motif within its vWFa domain. We also show that the MIDAS is important in HC1-mediated inhibition of the alternative pathway C3 convertase, via its interaction with C3, and demonstrate that HC1 can interact with vitronectin and other integrin ligands (e.g. small latent complexes of TGFβ) in a MIDAS-independent manner.

## Results

### Structure of rHC1

Our construct for recombinant HC1 (rHC1) encompasses the entire 638-residue mature protein sequence of human HC1 as defined by amino acid residues 35-672 in UniProt (ITH1, isoform A); see Supplementary Figure S1. Given that we observed weak metal ion-dependent dimerisation of rHC1 (see later), we conducted crystallisation screens using the D298A mutant that does not dimerise. Once crystals had been obtained with D298A, we were able to crystallise the wild type (WT) protein in similar conditions. The asymmetric unit of the crystals for both WT and D298A contained 2 independent copies of rHC1. Our crystal structure of rHC1 (at 2.34Å and 2.20Å resolution for WT (PDB: 6FPY) and D298A (PDB: 6FPZ), respectively; see Table 1) reveals that heavy chains are composed of 3 distinct domains (Figure 1). Its vWFa domain (residues 288-477) is inserted into a loop in an integrin-like hybrid domain (termed here HC-Hybrid1) composed of residues 266-287 and 478-543 (and linked by a disulphide bond (Olsen et al., 1998)). These two domains sit atop a large, novel, 16-stranded β-sandwich, composed of residues 45-265 and 601-652, which we call the HC-Hybrid2 domain (Figure 1A). The C-terminal end of the HC-Hybrid1 domain is connected to the final loops of the HC-Hybrid2 domain by 3 α-helices (residues 544-600). A construct-derived hexa-His tag (<u>AHHHHHHVGTGSNDDDDDKSP</u>), and residues 35-44, 631-636 and 653-672 of HC1, clearly present in the protein preparation as determined by mass spectrometry, were not visible in the electron density and are therefore assumed to be unstructured or highly conformationally labile. This includes the native C-terminus of HC1, which is covalently attached to CS in IαI and to HA in the context of HC•HA complexes. These missing residues were modelled using Small Angle X-ray Scattering (SAXS) data for (monomeric) D298A as a restraint target (Figure 1C,D); as can be seen, the AllosMod model fits better than the crystal structure alone to the experimental SAXS curve, with χ values of 1.56 and 2.68, respectively.

**Table 1.** Data collection and refinement statistics for rHC1.

|  | **WT – 6FPY** | **D298A – 6FPZ** |
| --- | --- | --- |
| **Wavelength** | 0.92Å | 0.92Å |
| **Resolution range** | 71 - 2.3 (2.4 - 2.3)<sup>a</sup> | 56.8 - 2.2 (2.3 - 2.2) |
| **Space group** | P 42 | P 42 |
| **Unit cell** | 158.8 158.8 65.4 90 90 90 | 159.7 159.7 65.79 90 90 90 |
| **Total reflections** | 455794 (45882) | 312045 (26776) |
| **Unique reflections** | 69109 (6862) | 83837 (8233) |
| **Multiplicity** | 6.6 (6.7) | 3.7 (3.3) |
| **Completeness (%)** | 91.94 (87.58) | 94.27 (88.13) |
| **Mean I/sigma(I)** | 8.02 (1.88) | 10.93 (2.05) |
| **Wilson B-factor** | 37.46 | 34.70 |
| **R-merge** | 0.1432 (0.8002) | 0.06619 (0.4731) |
| **R-meas** | 0.1554 (0.8681) | 0.07734 (0.5596) |
| **R-pim** | 0.05991 (0.3337) | 0.03928 (0.2917) |
| **CC1/2** | 0.994 (0.523) | 0.995 (0.478) |
| **CC\*** | 0.998 (0.829) | 0.999 (0.804) |
| **Reflections used in refinement** | 63610 (6027) | 79667 (7405) |
| **Reflections used for R-free** | 3194 (306) | 3983 (347) |
| **R-work** | 0.2317 (0.3810) | 0.2174 (0.3701) |
| **R-free** | 0.2605 (0.4195) | 0.2475 (0.3641) |
| **CC(work)** | 0.953 (0.726) | 0.954 (0.730) |
| **CC(free)** | 0.946 (0.696) | 0.948 (0.768) |
| **Number of non-hydrogen atoms** | 9702 | 10106 |
| **macromolecules** | 9319 | 9335 |
| **ligands** | 38 | 58 |
| **solvent** | 345 | 713 |
| **Protein residues** | 1201 | 1196 |
| **RMS(bonds)** | 0.004 | 0.003 |
| **RMS(angles)** | 1.03 | 0.92 |
| **Ramachandran favoured (%)** | 97.65 | 98.15 |
| **Ramachandran allowed (%)** | 2.27 | 1.85 |
| **Ramachandran outliers (%)** | 0.08 | 0.00 |
| **Rotamer outliers (%)** | 1.20 | 1.00 |
| **Clashscore** | 1.78 | 1.71 |
| **Average B-factor** | 49.67 | 45.75 |
| **macromolecules** | 49.92 | 45.76 |
| **ligands** | 62.82 | 71.64 |
| **solvent** | 41.55 | 43.54 |
| **Number of TLS groups** | 8 | 6 |

1 <sup>a</sup>Statistics for the highest-resolution shell are shown in parentheses.

**Figure 1:**
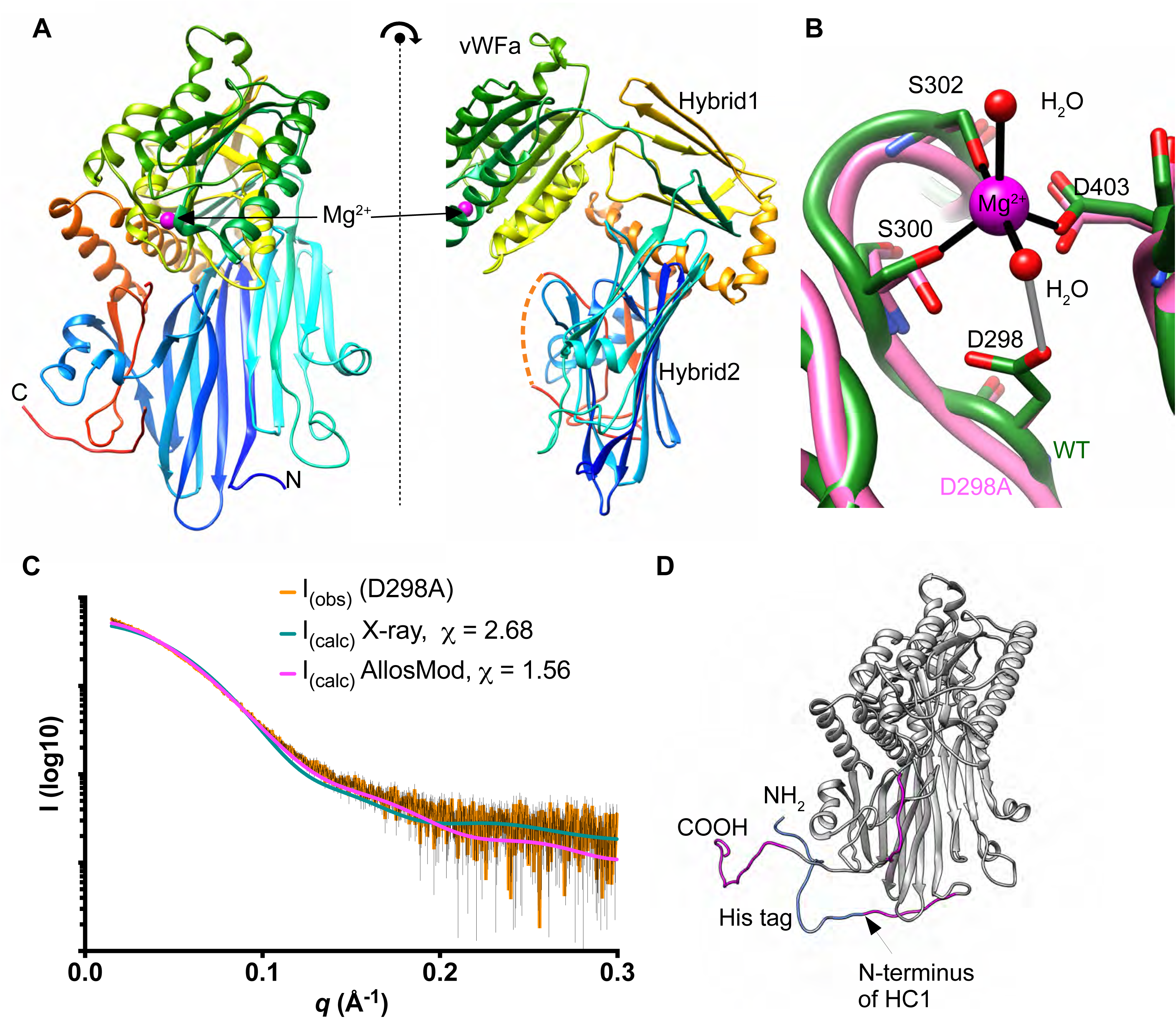
The crystal structure of HC1. **A)** Orthogonal views of the structure of rHC1, coloured from N-(blue) to C-(red) terminus, where domains and the bound Mg^2+^ ion are labelled; the dotted red line denotes resides 631-636, which are not visible in the crystal structure. **B)** Close up of the MIDAS site, showing metal co-ordination (black) and an important hydrogen bond (grey). The WT structure is shown in green and D298A structure (which lacks a Mg^2+^ ion) in pink. **C)** Raw SAXS data (orange with black error bars) of a rHC1 monomer (D298A), and back-calculated scattering curves based on the crystal structure of rHC1 alone or the crystal structure with the unstructured/flexible regions modelled in using Allosmod. **D)** Allosmod model of rHC1 with the N-terminal histidine tag (blue) and residues 35-44, 631-636 and 653-672 (pink) modelled based on SAXS restraints.

Despite low sequence identities (17% and 15%, respectively), the HC1 vWFa domain is structurally most similar to the vWFa domains from capillary morphogenesis protein 2 (CMG2; Lacy et al., 2004) and tumour endothelial marker 8 (TEM8; Fu et al., 2010), with PDBeFold Q-scores of 0.56 and 0.52 respectively. These are both transmembrane proteins that serve as functional receptors for the anthrax toxin (Liu et al., 2012). HC1 also shows significant structural similarity to the vWFa domains of various integrin I-domains, with the highest Q-score (0.50) for integrin α_M_ (ITGAM; also known as CD11b and as complement receptor type 3 (CR3)), and the vWFa domain of complement factor B (FB; Q-score 0.37); FB and ITGAM are C3-binding proteins, with roles in complement activation/amplification and complement-mediated phagocytosis, respectively (Ricklin et al., 2016). From the structure of WT rHC1 it is apparent that its vWFa domain contains a metal ion-dependent adhesion site (MIDAS) motif (Figure 1B), which was predicted from its sequence (Rugg et al., 2005); residues Asp298, Ser300, Ser302 and Asp403 chelate a magnesium ion, the identity of which can be inferred from the trigonal bipyramid co-ordination geometry, bond distances and refined atomic displacement parameters (ADPs). The D298A mutant lacks the Asp298 sidechain and has no bound Mg^2+^ ion but is otherwise very similar to the WT structure, with a RMSD between the two most similar chains of 0.24Å over 598 C-alpha atoms.

The HC-Hybrid1 domain of HC1 is composed of two 4-stranded β-sheets, where two of the β-strands are formed from amino acid residues before the vWFa domain and the remaining 6 from sequence after it; these regions are connected by a disulphide bond between Cys268 and Cys540. This arrangement of the HC-Hybrid1 and vWFa domains is reminiscent of integrin β-chains, as illustrated in Figure 2A,B for a comparison of rHC1 with ITGB3. Here the topologies of the β-strands are similar and, when the vWFa domains of HC1 and ITGB3 are superimposed, the ‘hybrid’ domains are ∼40° out of alignment (Figure 2C).

**Figure 2:**
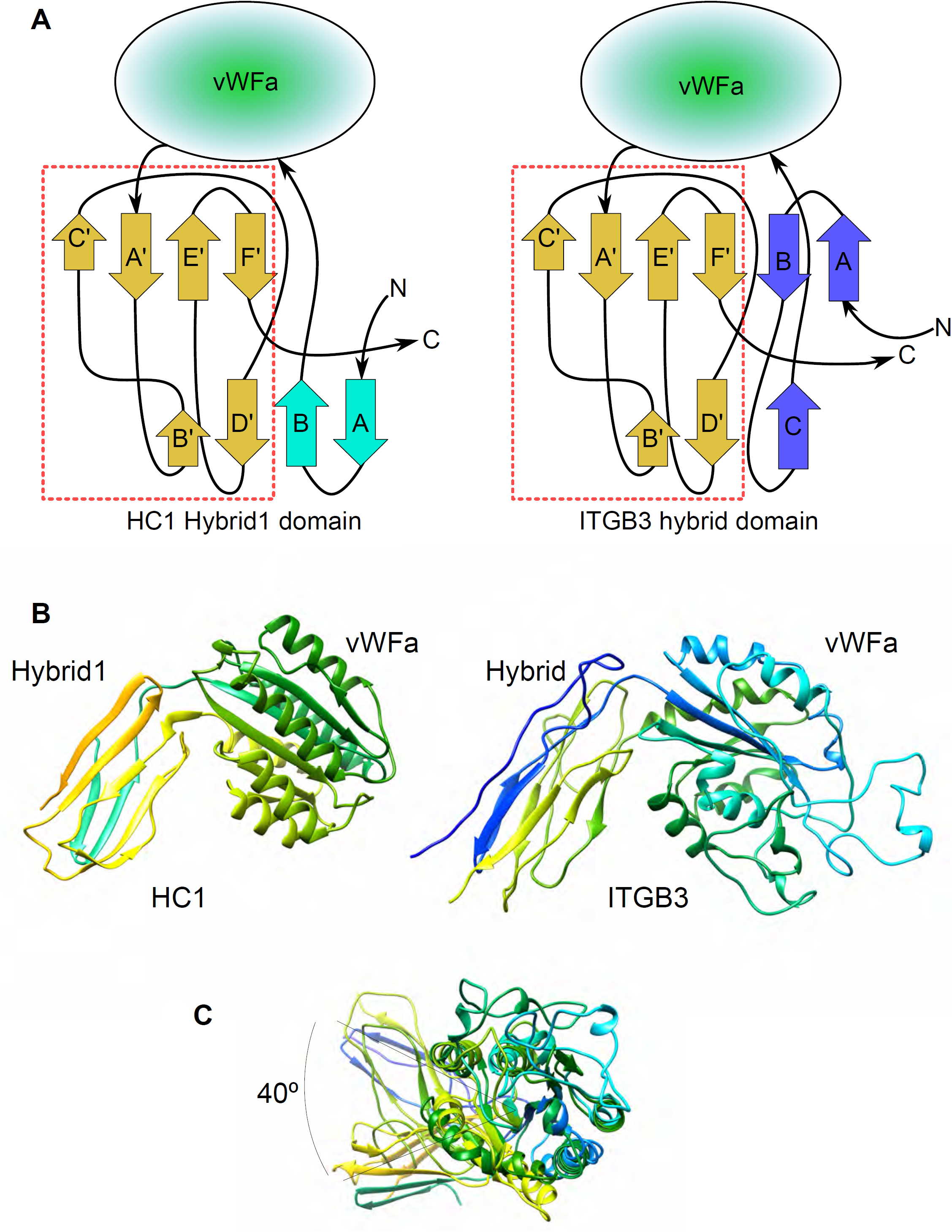
Integrin-like arrangement of vWFa and Hydrid1 domains in HC1 structure. **A)** Topologies of the HC-Hybrid1 domain from HC1 (left) and the hybrid domain from human integrin β3 chain (ITGB3) (right); the arrangements of β-strands in the sequences following the vWFa domains are essentially identical (dashed red box). **B)** Side-by-side views of the ‘hybrid’ and vWFa domain pairs of HC1 (left) and ITGB3 (right). **C)** The ‘hybrid’ domains are displaced by ∼40° when the vWFa domains of HC1 and ITGB3 are superimposed.

The HC-hybrid1 domain is a variant of the fibronectin type-III (FnIII) fold, and its closest structural match in mammalian extracellular proteins is the third FnIII domain from integrin IGTB4 (Alonso-García et al., 2015), with a RMSD between the structures of 1.55Å. Curiously, the closest structural similarity overall is to the EAR domain of gamma 2 adaptin (RMSD – 1.28Å), a protein found in the clathrin adaptor complex, which is involved in intracellular protein transport and is hijacked in hepatitis B infection (Jürgens et al., 2013).

The HC1-Hybrid2 domain has a unique structure composed of 16 β-strands arranged into two β-sheets. It is most similar to a “domain of unknown function” from PDB 4G2A, for which there is no accompanying publication. In the HC1-Hybrid2 domain one β-sheet is continuous, but the other has a missing strand, with a break in the hydrophobic core of the domain. The resulting subdomains are structurally homologous to immunoglobulin-like carbohydrate binding domains (residues 44-170) and a jelly-roll fold (residues 173-263).

### rHC1 forms MIDAS and metal-ion dependent dimers

During preparative size exclusion chromatography (SEC) of rHC1 for crystallisation the presence of a small amount of dimer was observed when we included metal ions in the buffer. Given that HC-HC interactions have been proposed to non-covalently cross-link HC•HA complexes (Yingsung et al., 2003), we explored this phenomenon further with both analytical ultracentrifugation (AUC) and SAXS. Velocity AUC revealed that in the presence of magnesium, WT rHC1, while mostly monomeric, formed dimers (Figure 3A); here sedimentation coefficients (s_(20,w)_) of 4.59 S and 6.11 S were obtained. Equilibrium AUC conducted at a range of magnesium ion concentrations (Supplementary Figure S2) showed that this interaction was indeed Mg^2+^ dependent, although the affinity is rather weak (*K*_D_ = 35.3µM at 1mM MgCl_2_; Figure 3B). We used high-throughput SAXS screening, generating D_max_ (maximum dimension) values, as an efficient way of determining dimerisation in a range of different metal ion conditions (Table 2). Magnesium chloride and manganese chloride supported dimerisation, whereas calcium chloride and EDTA did not. D298A did not form Mg^2+^- or Mn^2+^-dependent dimers and, therefore, we concluded that the dimerisation activity of rHC1 requires a correctly formed and metal ion-occupied MIDAS that can accommodate either a magnesium or manganese ion.

**Table 2.** Biophysical analysis of rHC1 dimerisation. Radius of gyration (R_g_), maximum dimension (D_max_), approximate molecular weight (Mwt) and sedimentation coefficient (s_w(20,w)_) values were derived from SAXS and AUC data for WT and D298A rHC1. All D298A data, and WT data collected in the presence of 2.5mM EDTA, are consistent with a monomeric state. WT rHC1 with 5mM MgCl_2_ or 5mM MnCl_2_ is dimeric. Data from “As purified” WT rHC1 and in 5mM CaCl_2_ are consistent with a mixture of monomer and dimer; this is presumably due to trace amounts of Mg^2+^ ions present in various buffer components. AUC data are derived from equilibrium experiments performed in triplicate at 3 different speeds; SAXS data are from data processed by AUTORG and with DATGNOM (i.e. with no imposed constraints). The molecular weight of an rHC1 monomer from intact mass spectrometry is 73,802 Da.

| Protein | Metal ion | $R_g^{SAXS}$<br>(Å) | $D_{max}^{SAXS}$<br>(Å) | Mwt<sup>SAXS</sup><br>(kDa)<sup>b</sup><br>(Ratio SAXS<br>mass/Intact mass) | $s_{(20,w)}^{AUC}$<br>(S) | $s_{(20,w)}^{SAXS}$<br>(S) |
| --- | --- | --- | --- | --- | --- | --- |
| rHC1 WT | None<sup>a</sup> | 35.2 | 123 |  |  |  |
|  | EDTA<br>(2.5mM) | 31.8 | 112 | 78<br>(1.06) | 4.39 | 4.50 |
| | $Mg^{2+}$ (5mM) | 49.5 | 170 | 140<br>(1.89) | 4.59(monomer)<br>6.11(dimer) | 6.24 |
| | $Mn^{2+}$ (5mM) | 51.2 | 172 | | | |
| | $Ca^{2+}$ (5mM) | 38.5 | 135 | | | |
| rHC1 D298A | None<sup>a</sup> | 33.3 | 114 |  |  |  |
| | $Mg^{2+}$ (5mM) | 33.3 | 114 | | | |
| | $Mn^{2+}$ (5mM) | 32.8 | 105 | | | |
| | $Ca^{2+}$ (5mM) | 32.8 | 115 | | | |
| IαI | $Mg^{2+}$ (5mM) | 49.2 | 170 | | | |

<sup>a</sup>As purified
<sup>b</sup>Calculated using the method of Rambo & Tainer (2013) *Nature* **496**, 477-481

**Figure 3:**
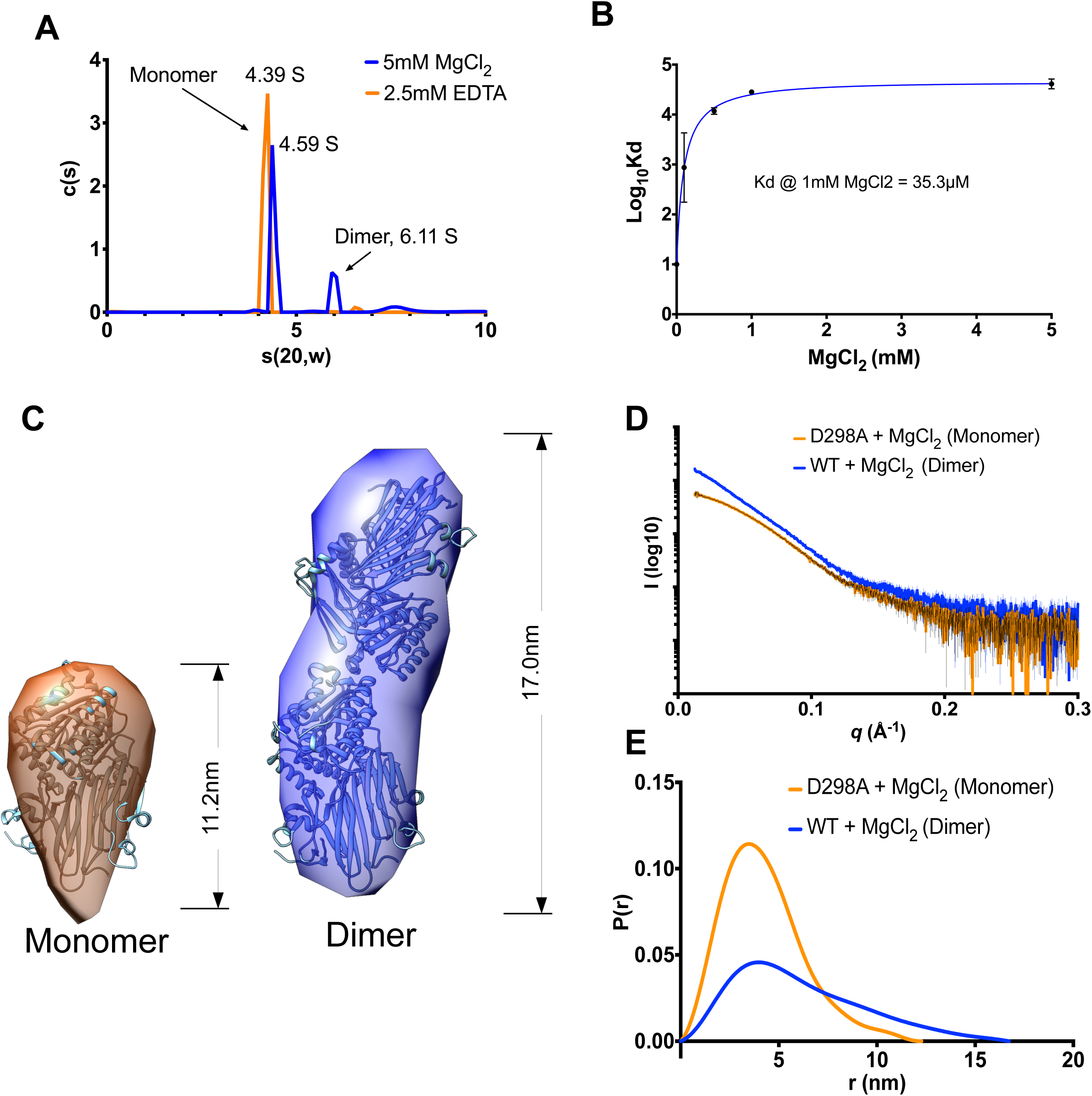
HC1 forms metal ion-dependent dimers. **A)** A plot of sedimentation coefficient distributions (c(s)) vs s(apparent)) for WT rHC1 derived from velocity AUC analysis. In the presence of 2.5mM EDTA (orange), 93% of the rHC1 protein is in a monomeric state (s_(20,w)_ = 4.39 S) and there is no detectable dimer present; in 5mM MgCl_2_ (blue) 64% of the protein is monomeric (s_(20,w)_ = 4.59 S) and 21% of material is dimeric (s_(20,w)_ = 6.11 S). **B)** Plot of Log_10_*K*_D_ vs MgCl_2_ concentration, derived from equilibrium AUC measurements. At 0mM MgCl_2_ (achieved by conducting the experiment in 2.5mM EDTA) no dimerisation was detected. Maximal binding affinity (for self-association of the rHC1 dimer) was reached at ∼1mM MgCl_2_, i.e. close to the concentration of free Mg^2+^ ions in plasma. **C)** *Ab initio* SAXS models for the HC1 monomer (left) and dimer (right) where the HC1 structure has been modelled into the SAXS envelopes. **D)** Buffer-subtracted SAXS scattering curves for HC1 D298 monomer (orange) and WT dimer (blue) and **E)** their derived P(r) vs Distance plots, consistent with WT HC1 forming an elongated Mg^2+^-dependent dimer and the MIDAS site mutant (D298A) being monomeric.

SAXS data for the rHC1 monomer (D298A in MgCl_2_) and dimer (wild type in MgCl_2_) were used to obtain low resolution *ab initio* solution structures (Figure 3C-E); see Supplementary Figure S3 for analysis of SAXS data, showing the rHC1 monomer and dimers to be folded and rigid. For the monomer it was apparent that the crystal structure for rHC1 could be well accommodated within the SAXS envelope. On the other hand, when two HC1 molecules were fitted into the envelope for the rHC1 dimer the fitting was ambiguous; the model presented in Figure 3C gives the best overall fit-to-map correlation, but other models give similar scores. One likely explanation is that a conformational change occurs in the HC1 structure on dimerisation. Moreover, small differences between the sedimentation coefficients determined by velocity AUC for the monomer species in EDTA (4.39 S) and MgCl_2_, (4.59 S) indicate that metal ion binding induces a structural change in the monomeric protein prior to dimer formation; i.e. consistent with a recent biochemical analysis (Scavenius et al., 2016). However, while this change is evident in the solution phase, we saw no such difference between the crystal structures for WT HC1 with a bound Mg^2+^ ion and the metal ion-free D298A mutant. This could be due to the fact that the initial crystallisation conditions were obtained from D298A protein, and that these conditions stabilise the protein in a monomeric configuration. The potential dimers observed in the crystal lattice do not correlate with the SAXS data.

### rHC1 structure enables modelling of inter-alpha-inhibitor

We recorded SAXS data for IαI purified from human plasma and found that even in the presence of 5mM MgCl_2_ it is monomeric (Figure 4A, B and Supplementary Figure S4); IαI, which is likely rigid, has an elongated shape (Figure 4C), with a D_max_ value of 17.0nm, which is similar to that for the HC1 dimer (Figure 3C, Table 2). The SAXS data (collected in HEPES buffered saline with 2mM MgCl_2_) were used to generate an *ab initio* solution structure for IαI and thereby determine the likely quaternary organisation of the IαI complex (Figure 4C), i.e. using the structures of bikunin (Xu et al., 1998) and rHC1, and a homology model of HC2 based on the HC1 coordinates (determined here) and experimentally determined disulphide bonds (Olsen et al., 1998). The three protein chains of IαI could be readily fitted within the SAXS envelop with the bikunin chain being accommodated in a small lobe at one end and the two HCs arranged asymmetrically in the larger lobe; this positioning would place the C-terminal peptides of HC1 and HC2 on the same face, making them close enough to take part in the observed CS conjugation. The IαI model shown in Figure 4C was used to back calculate SAXS data, where this was found to have reasonable agreement with the experimentally derived scattering data; i.e. a χ = 7.21 for I_(obs)_ vs I_(model)_.

**Figure 4:**
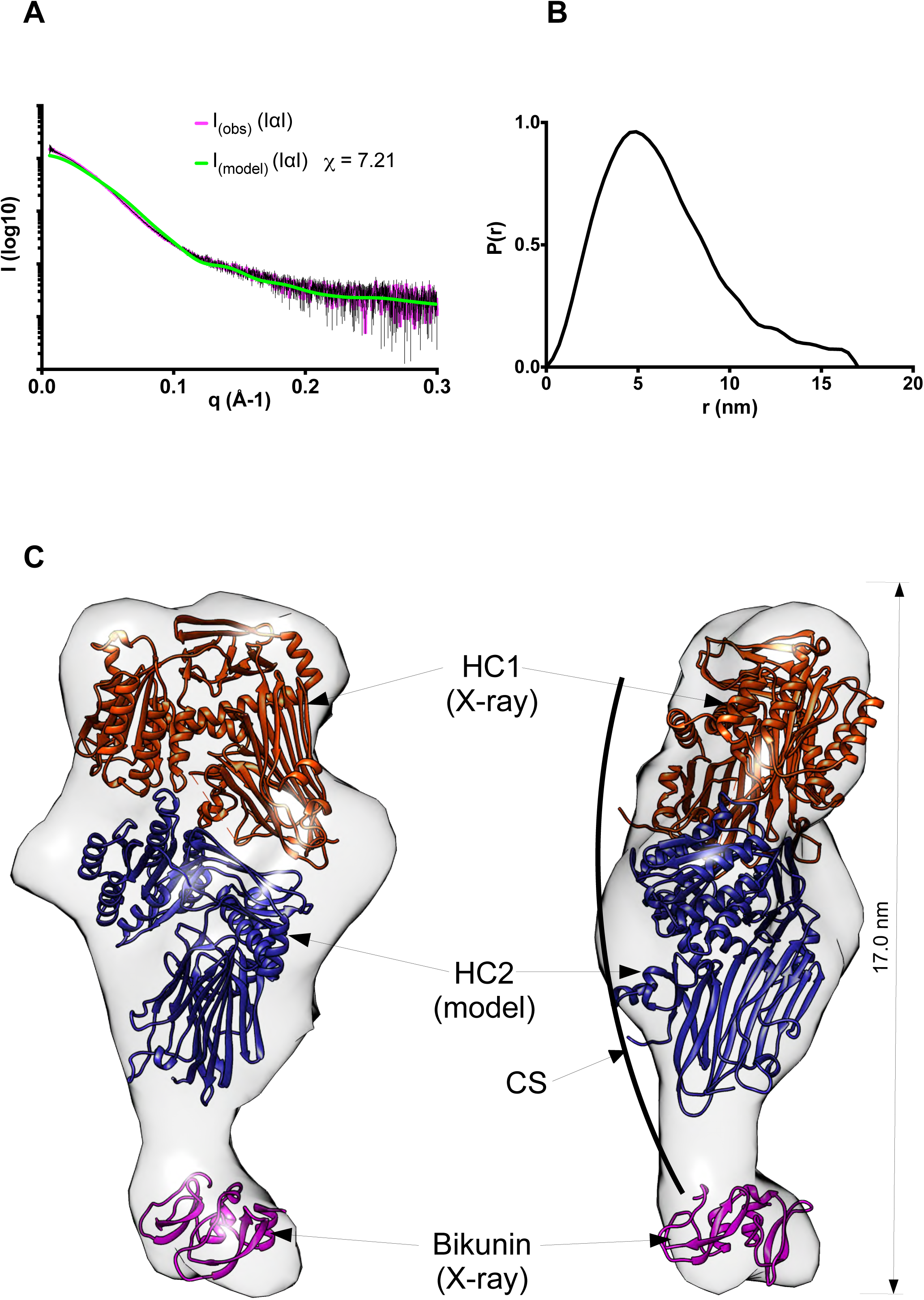
The quaternary structure of inter-α-inhibitor. **A)** Raw SAXS data for IαI (obs) in the presence of 2mM MgCl_2_ fitted to scattering data (pink with black error bars) derived from the pseudo atomic model (model) in **(C)** calculated using Allosmod-FoXs. **B)** P(r) vs Distance plot showing that IαI has an elongated and asymmetric shape. **C)** Orthogonal views of the SAXS envelope of IαI (transparent grey surface) determined *ab initio* from the SAXS scattering curve, with structures of bikunin (PDB 1BIK; pink) and rHC1 (determined here; brown) and a threading model of HC2 (based upon the structure of HC1; blue), modelled in. The CS chain is shown schematically to indicate its expected position relative to the three protein chains.

### rHC1 inhibits the alternative complement pathway in a MIDAS-dependent manner

Given there is evidence that IαI binds to C3 and that heavy chains are inhibitors of the alternative and classical complement pathways (Okroj et al., 2012), we investigated whether rHC1 could interact with C3 (i.e. the central component of complement). Initial buffer screening using surface plasmon resonance (SPR), revealed that rHC1 interacted with C3 in a Mn^2+^-ion-dependent manner, which is mediated via the HC1 MIDAS motif since the D298A mutant exhibited no binding activity (Figure 5A; Table 3); there was also an interaction in Mg^2+^ (albeit of lower apparent affinity) but there was no binding in the presence of Ca^2+^ or EDTA (data not shown). Full SPR analysis (in 2 mM MgCl_2_/2 mM MnCl_2_) determined that the *K*_*D*_ for the rHC1-C3 interaction was ∼360 nM, i.e. a similar affinity to the binding of C3 to IαI (*K*_D_ = ∼660nM; Table 3). When we tested rHC1 in a functional assay of complement activation we found that the WT protein, but not the D298A mutant (data not shown), was able to dose-dependently inhibit the activity of the alternative pathway C3 convertase (C3bBb) with an IC_50_ of 980 nM (Figure 5B).

**Table 3.** Analysis of rHC1-ligand interactions by SPR.

| Immobilised ligand | Analyte | Buffer conditions | Replicates | $K_D$ (nM) ± SD |
| --- | --- | --- | --- | --- |
| C3 | rHC1 (WT) | 2mM $Mg^{2+}$/2mM $Mn^{2+}$ | 3 | $364 \pm 78$ |
| C3 | rHC1 (WT) | 2mM $Mn^{2+}$ | 3 | $473 \pm 329$ |
| C3 | rHC1 (D298A) | 2mM $Mg^{2+}$/2mM $Mn^{2+}$ | 3 | NB<sup>a</sup> |
| C3 | IαI | 2mM $Mg^{2+}$/2mM $Mn^{2+}$ | 3 | $659 \pm 205$ |
| rHC1 (WT) | Vitronectin | 10mM EDTA | 3 | $0.192 \pm 0.014$ |
| rHC1 (WT) | Vitronectin | 2mM $Mg^{2+}$/2mM $Mn^{2+}$ | 3 | $0.118 \pm 0.011$ |
| rHC1 (D298A) | Vitronectin | 10mM EDTA | 3 | $0.138 \pm 0.040$ |
| rHC1 (D298A) | Vitronectin | 2mM $Mg^{2+}$/2mM $Mn^{2+}$ | 3 | $0.096 \pm 0.044$ |
| rHC1 (ΔvWFa) | Vitronectin | 10mM EDTA | 3 | $0.097 \pm 0.049$ |
| rHC1 (WT) | TGFβ1-LAP | 2mM $Mg^{2+}$/2mM $Mn^{2+}$ | 3 | $16.1 \pm 5.5$ |
| rHC1 (D298A) | TGFβ1-LAP | 2mM $Mg^{2+}$/2mM $Mn^{2+}$ | 3 | $14.9 \pm 2.7$ |
| rHC1 (WT) | TGFβ2-LAP | 2mM $Mg^{2+}$/2mM $Mn^{2+}$ | 3 | $8.8 \pm 0.6$ |
| rHC1 (D298A) | TGFβ2-LAP | 2mM $Mg^{2+}$/2mM $Mn^{2+}$ | 3 | $12.9 \pm 1.7$ |
| rHC1 (WT) | TGFβ3-LAP | 2mM $Mg^{2+}$/2mM $Mn^{2+}$ | 3 | $9.9 \pm 1.4$ |
| rHC1 (D298A) | TGFβ3-LAP | 2mM $Mg^{2+}$/2mM $Mn^{2+}$ | 3 | $3.1 \pm 0.8$ |
| rHC1 (WT) | LAP (TGFβ1) | 10mM EDTA | 3 | $2.0 \pm 0.3$ |
| rHC1 (WT) | TGFβ1 | 10mM EDTA | 3 | NB |
| rHC1 (WT) | TGFβ3 | 10mM EDTA | 3 | NB |
| rHC1 (WT) | LTBP1 NT1 | 10mM EDTA | 3 | $5.1 \pm 0.5$ |
| rHC1 (WT) | LTBP1 NT1 | 2mM $Mg^{2+}$/2mM $Mn^{2+}$ | 3 | $4.6 \pm 0.2$ |
| rHC1 (WT) | LTBP1 EGF | 2mM $Mg^{2+}$/2mM $Mn^{2+}$ | 3 | NB |
| rHC1 (WT) | LTBP1 CT | 2mM $Mg^{2+}$/2mM $Mn^{2+}$ | 3 | NB |

<sup>a</sup>NB = no binding

**Figure 5:**
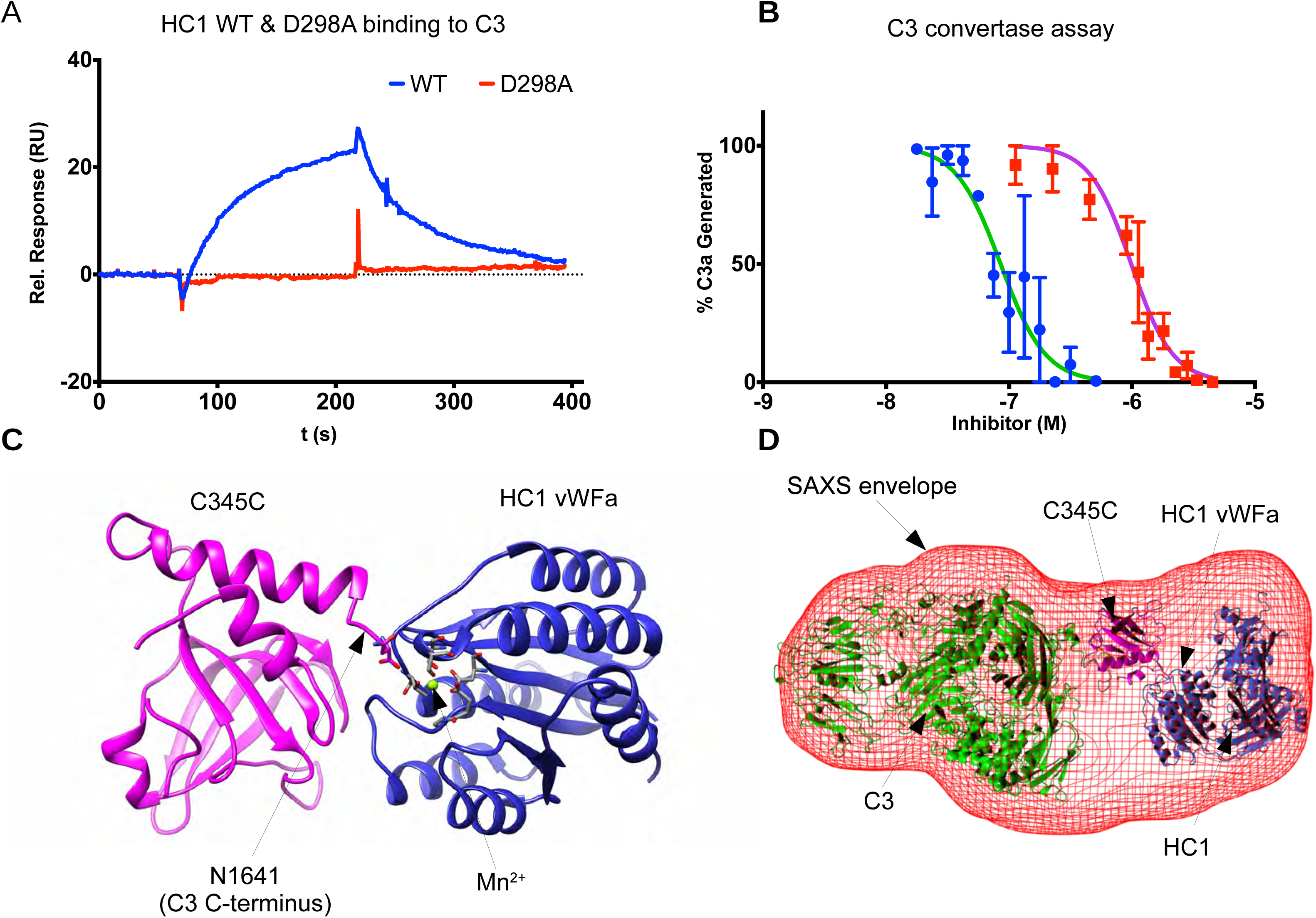
HC1 inhibits the alternative pathway C3 convertase activity through interaction with C3. **A)** SPR analysis for the interaction of rHC1 (WT and D298A) with C3 (in 2mM MnCl_2_), where the lack of binding of the D298A mutant indicates an essential role for the MIDAS site. **B)** rHC1 proteins (WT and D298A) were compared with factor H (FH) in an alternative pathway C3 convertase assay (containing Mg^2+^ and Mn^2+^ ions), where the proteolytic release of C3a was quantified (by SDS-PAGE) as a surrogate for the conversion of C3 into C3b. Only WT rHC1 had inhibitory activity; data for D298A, not shown. Mean values (± SD) were derived from independent experiments performed in triplicate. Data were fitted using Graphpad Prism to derive IC_50_ values for rHC1 and FH control. **C)** An *in silico* model of the C3 C-terminal C345C domain (pink) bound to the vWFa domain of HC1 (blue). Here a Mn^2+^ ion (green) occupies the MIDAS of HC1 (with co-ordinating residues shown in stick representation) and co-chelates the carboxy-terminal amino acid (Asn) of C3b. **D)** An *ab initio* SAXS structure was determined for the rHC1-C3 complex (red mesh), where C3 and HC1 molecules, interacting as in **(C)**, could be accommodated.

The divalent cation and MIDAS-dependent interaction of rHC1 with C3 is reminiscent of the manner in which FB associates with C3b (activated C3) to form the C3 convertase; this is mediated by the vWFa domain in FB binding to the C-terminus of C3/C3b via co-chelation of a Mg^2+^ ion bound to FB’s MIDAS motif (Forneris et al., 2010). *In sillico* modelling of the HC1 vWFa and the C-terminal domain (C345C) of C3/C3b (Figure 5C) reveals that a MIDAS-mediated interaction is indeed feasible and consistent with a low resolution SAXS structure determined for the rHC1-C3 complex (Figure 5D and Supplementary Figure S5); while the complex is folded and globular, Porod-Debye analysis indicated that it had some flexibility.

### HC1 binds to integrin ligands in a MIDAS- and vWFa-independent manner

Given the structural similarity of HC1 to integrin β-subunits and our finding that rHC1 dimerises and binds to complement C3 in a metal ion-, and likely MIDAS motif-, dependent manner we wanted to further explore its interaction with integrin ligands; as described above, IαI is known to bind to vitronectin, where the vWFa domain has been implicated in binding (Adair et al., 2009). Our SPR analysis (Supplementary Figure S6) showed that rHC1 binds with high affinity to vitronectin, however, to our surprise this was independent of metal ions (Table 3); essentially identical shaped binding curves were seen for experiments in Mg^2+^/Mn^2+^ and EDTA (data not shown). Moreover, the D298A mutant and a construct where the vWFa domain had been removed (ΔvWFa) both bound to vitronectin with very similar affinities to the WT protein (Supplementary Figure S6C; Table 3).

We also investigated the binding of rHC1 to the small-latent complexes (SLC) of TGFβ1, 2, and 3, in which the growth factors are coupled to latency-associated peptides (LAP); TGFβ1-LAP and TGFβ3-LAP (which both contain an RGD motif) are activated by the α_V_β_6_ and α_V_β_8_ integrins, in response to mechanical stress, in a metal ion- and MIDAS-dependent manner (Annes et al., 2004; Shi et al., 2011; Worthington et al., 2015). As shown in Supplementary Figure S6D, we found that rHC1 could interact tightly with TGFβ1-LAP, TGFβ2-LAP and TGFβ3-LAP, where the affinity (*K*_D_ ∼10 nM) was essentially identical for the WT and D298A mutant (Table 3); moreover, similar binding was seen in EDTA (not shown). Together this demonstrates that the interactions are independent of metal ions and do not involve HC1’s MIDAS motif. Additional SPR experiments revealed that rHC1 interacts with the LAP peptide (analysed for the LAP from TGFβ1; *K*_*D*_ = 2 nM) but did not bind to the mature growth factor (i.e. TGFβ1 and TGFβ3). SLCs associate with latent TGFβ binding proteins (LTBP) to form large latent complexes (LLCs) (Robertson et al., 2015); this mediates matrix sequestration and regulates the activation of latent TGFβ. We tested whether rHC1 could bind to LTBP1 and found that it interacts with the N-terminal region (NT1), again in a metal ion-independent manner, but not with the C-terminal (CT) or EGF regions (Table 3).

## Discussion

Here we have determined the first crystal structure for a heavy chain of the IαI/ITIH family. Given the similarity of the prototypical HC1 to the 5 other HC proteins encoded in the human genome (32-54% sequence identity), our study defines the canonical structure for a heavy chain, allowing the modelling of other family members. In this regard, we generated a homology model of HC2 that, along with the structure for rHC1 (and bikunin), allowed us to infer the quaternary organisation of IαI itself. Our SAXS-based modelling of IαI (Figure 4) reveals that this unusual CS proteoglycan forms an elongated structure, but with a compact arrangement of the 3 protein chains as also inferred in a recent study (Scavenius et al., 2016).

Unlike IαI, which is monomeric, rHC1 forms a dimer in solution. Given the metal ion-dependence of dimer formation (requiring Mg^2+^ or Mn^2+^; Table 2) and the lack of dimerisation by the D298A mutant, the MIDAS motif within the vWFa domain clearly plays an essential role in mediating this protein-protein interaction. It is possible that an Asp or Glu sidechain on one HC1 monomer could engage with the metal ion within the MIDAS of the other HC1; e.g. to effect a conformational change, thereby altering the orientation of the vWFa domain relative to the rest of the protein and leading to the dimer dimensions indicated by SAXS (Figure 3C). This is reminiscent of how metal ion and ligand occupancy of an integrin MIDAS can transduce a conformational change that causes the hybrid and vWFA domains to swing away from one another during integrin activation (Luo et al., 2007; Wang et al., 2017). The arrangement of the HC1 and HC2 vWFa domains in our IαI model (Figure 4C) indicate that such interactions would be sterically precluded, explaining why IαI does not dimerise.

It is well established that HC1, HC2 and HC3 can become covalently attached to the polysaccharide HA, via transesterification reactions catalysed by TSG-6, e.g. in the context of ovulation and inflammation; see (Day and Milner, 2019). This reaction requires the presence of Ca^2+^ and Mg^2+^/Mn^2+^ ions (Briggs et al., 2015) and occurs via the formation of covalent TSG-6•HC intermediates (Rugg et al., 2005). There is a Ca^2+^ ion-binding site in TSG-6, which we have shown previously to be essential for TSG-6•HC formation (Briggs et al., 2015), and our finding here that a Mg^2+^or Mn^2+^ ion can be accommodated within the vWFa domain of HC1 (Figure 1) provides strong evidence that HCs are the source of these metal ions. Moreover, solving the heavy chain structure will facilitate refinements in our understanding of the mechanisms underlying the transfer of HCs onto HA.

The dimerization of rHC1 provides the first direct evidence that homotypic HC-HC interactions might contribute to the stabilisation of HC•HA-rich matrices. Given that the C-terminal 19 amino acid residues of HC1 (which were not visible in the crystal structure) are likely to form a flexible linker, this protein-protein interaction is unlikely to be affected by whether the C-terminus of HC1 is covalently attached to HA or not. We found that the HC1-HC1 interaction is rather weak (*K*_d_ ∼40 μM at physiological Mg^2+^ concentrations; Figure 3B) indicating that, for this heavy chain, at least, binding is likely to be highly transient. As yet we do not know whether other HCs self-associate in this way or indeed the nature/affinities of heterotypic HC-HC interactions. However, it seems reasonable to propose that low affinity binding between HCs could mediate the aggregation of HC•HAs seen in synovial fluids from rheumatoid arthritis patients (Yingsung et al., 2003) and that this, combined with more stable interactions between HCs and PTX3 (Baranova et al., 2014), underpin the formation and crosslinking of the cumulus extracellular matrix during COC expansion. Furthermore, dynamic HC-HC interactions could make an important contribution to the mechanical properties of tissues; for example, they might explain the elasticity and extreme softest of the cumulus matrix (Chen et al., 2016). Certainly HC•HAs have different compositions of heavy chains in different tissue contexts (Day and Milner, 2019) and it seems likely that this will engender distinct hydrodynamic and functional properties.

We found that rHC1 was able to bind to complement C3 with moderate affinity (*K*_D_ ∼360 nM; Table 3), thereby identifying this complement component as a novel heavy chain ligand. Modelling of the rHC1-C3 complex (Figure 5) demonstrated that this interaction could be mediated via the C-terminus of C3 co-chelating the metal ion within the MIDAS of HC1. This also provides a plausible mechanism by which rHC1 (and potentially IαI) inhibits the activity of the alternative pathway C3 convertase (Figure 5C), by acting as a competitor of the interaction between FB and C3. Displacement of FB may also explain how IαI inhibits the factor D-mediated cleavage of FB to Bb (Okroj et al., 2012), as this reaction requires FB to be associated with C3. In our functional assays, rHC1 was an approximately 10-fold weaker inhibitor compared to human factor H (FH), the only established negative regulator of the alternative pathway in the solution-phase (see (Parente et al., 2017)). While IαI and FH have similar concentrations in serum, in tissues where HC1 accumulates via covalent attachment to HA, its complement inhibitory activity could serve to dampen the innate immune response. HC1-mediated inhibition of complement activation might be particularly important during ovulation, where plasma proteins (including complement components and IαI) ingress into the ovarian follicle when the blood-follicle barrier breaks down, i.e. to provide protection to the COC prior to ovulation.

Our discovery that the vWFA of HC1 shares high structural similarity with those of TEM8 and CMG2 may be significant given that these proteins are known to be functional receptors for the anthrax toxin (Liu et al., 2012) and because IαI has been shown to protect against anthrax intoxication (Opal et al., 2011; Singh et al., 2010). The latter has previously been attributed to the activity of the bikunin chain in inhibiting furins/preprotein convertases, which are proteases that have a critical role in the assembly of the anthrax toxin protective antigen (Opal et al., 2005). The protective antigen binds to the host cell surface by utilising the receptors CMG2 and TEM8 (Liu et al., 2012), both of which contain vWFA domains that mediate the interaction via their MIDAS motifs (Fu et al., 2010; Lacy et al., 2004), in a similar manner to how integrins interact with their ligands. Thus, our data are consistent with a mechanism whereby IαI and HCs act as decoy receptors for the anthrax toxin and sequester the toxin in the fluid phase, preventing it from binding to membranes and forming the pores that give rise to the toxin’s cytotoxic activity.

We have identified that rHC1 binds to vitronectin (a ligand of the α_V_β_6_ integrin) with very high affinity (*K*_D_ ∼0.2 nM; Table 3), consistent with a previous report (Adair et al., 2009). However, our data clearly demonstrate for HC1 (at least) that the interaction with vitronectin does not involve the vWFa domain and is thus not a typical RGD-mediated MIDAS co-chelation interaction; e.g. an rHC1 construct lacking the entire vWFa domain bound to vitronectin with similar affinity to the wildtype protein (Supplementary Figure S6C; Table 3). The finding that the binding of IαI to vitronectin is inhibited by RGD peptides (Adair et al., 2009) is intriguing and suggests that even though this interaction is not mediated by metal ions the integrin-binding site in vitronectin may be involved. Further work is needed to investigate this possibility and determine the effect of HC1 on α_V_β_6_-vitronectin interactions.

In light of the tight but non-canonical interaction of rHC1 with vitronectin, and given that TGFβ1 and TGFβ3 interact with α_V_β_6_ via RGD sequences within their latency-associated peptides (Annes et al., 2004; Shi et al., 2011), we screened the 3 small latent complexes of TGFβ, for binding to rHC1. We found that all three SLCs interacted with rHC1 with high affinity (*K*_D_ ∼10 nM; Table 3), including TGFβ2-LAP that doesn’t have an RGD motif. As in the case of vitronectin, the D298A mutant of rHC1 (with a defective MIDAS) bound the SLCs with similar affinities to WT rHC1. Additional SPR data indicated that rHC1 binds to the LAP rather than the mature growth factors and also interacts with the N-terminal region of LTBP1, which associates with TGFβ-LAP to form the large latent complex (LLC). Given that the LLCs sequester TGFβs in the matrix (Robertson et al., 2015), through interactions with both the N- and C-terminal regions of LTBP1, it seems reasonable to suggest that HC1 may play a role in regulating the bioavailability of these important growth factors/cytokines. In this regard, whether HC1 acts in an analogous fashion to α_V_β6, i.e. to mechanically activate the release of mature TGFβ (Buscemi et al., 2011), or whether it stabilises the LLC remains to be determined. The latter seems more likely based on its binding to both LAP and LTBP1 and is consistent with the finding that HC•HA complexes present in the human amniotic membrane, which are reported to only contain HC1 (Zhang et al., 2012), have been found to be potently tissue protective with anti-fibrotic activity (Ogawa et al., 2017).

In summary, this study has identified that HC1 has a structural organisation reminiscent of an integrin β-chain, including vWFa/hybrid domains and a functional MIDAS motif that mediates some but not all of its ligand-binding interactions. Our novel findings that HC1 can inhibit the complement system and has the potential to modulate TGFβ activity indicates that this protein is likely to be an important regulator of the innate and adaptive immune systems, for example, when it becomes covalently associated in the extracellular matrix during inflammation.

## Materials and Methods

### Protein production

The rHC1 proteins (WT, D298A and a ΔvWFa mutant, lacking residues 288-478) and the recombinant domains of LTBP1 (NT1, EGF and CT) were expressed and purified as described previously (Baranova et al., 2013; Troilo et al., 2016). LAP (from TGFβ1), TGFβ1, TGFβ1-LAP, TGFβ2-LAP, TGFβ3 and TGFβ3-LAP were obtained from R&D Systems, vitronectin was from PeproTech and complement C3 from Merck. IαI was purified from human plasma as described previously (Enghild et al., 1989).

### Crystallography of rHC1

WT rHC1 and the D298A mutant were crystallised by mixing 1μl of protein (10mg/ml in 10mM HEPES pH 7.5, 50mM NaCl) with an equal amount of crystallisation mother liquor (100mM HEPES, pH 7.5, 100mM sodium acetate, 10% (w/v) PEG8K, 20% (v/v) glycerol). Crystals appeared within one week. Native diffraction data were collected to 2.20Å (D298A) and 2.34Å (WT) and the data were indexed, integrated and scaled using DIALS (Waterman et al., 2016), POINTLESS (Evans, 2011), AIMLESS/SCALA (Evans et al., 2006; Evans et al., 2013) and cTRUNCATE (Evans, 2011) as implemented in the Xia2 pipeline (Winter et al., 2013). The data were phased using the SIRAS method and a K_2_PtCl_4_-derivatised D298A crystal. The substructure was solved, and the data phased, density modified and the chain partially traced using PHENIX AutoSol (Terwilliger et al., 2009). Both the WT and D298A models were rebuilt and refined to convergence using the COOT (Emsley et al., 2010) and PHENIX Refine (Afonine et al., 2012) packages. Data collection statistics are shown in Table 1. The refined models have been deposited in the PDB databank with accession codes 6FPY (WT) and 6PFZ (D298A).

### Small angle X-ray scattering and modelling of Iα I, rHC1 and rHC1-C3 complex

SAXS data were collected at beamline P12, PetraIII, DESY (Blanchet et al., 2015). Proteins (rHC1 or IαI) at 1.25, 2.5 and 5.0 mg/ml were prepared in HEPES buffered saline (pH 7.5). Data were reduced using PRIMUS/GNOM (Konarev et al., 2003; Svergun, 1992). The R_g_ and D_max_ values shown in Table 2 were calculated automatically using AUTORG and DATGNOM (Petoukhov et al., 2007) to prevent bias or subjective interpretation. *Ab initio* models were created using the DAMMIF/DAMMIN packages (Franke and Svergun, 2009; Svergun, 1999); 20 models were made using DAMMIF in slow mode. The averaged model from DAMMIF was refined to convergence using DAMMIN. Modelling of residues missing from the crystal structure was done using the Allosmod-FoXs server (Weinkam et al., 2012). Modelling of the dimeric form of HC1 was carried out as for the monomeric form, although P2 symmetry was enforced once it was determined that the data corresponded to a dimer. Rigid body docking of the HC1 structure into the resulting DAMMIN envelope was performed in UCSF Chimera for both monomeric and dimeric HC1. The resolution of the resulting map used for fitting was determined using SASRES (Tuukkanen et al., 2016); this was 43Å for the monomer and 64Å for the dimer. In modelling of the dimer, we enforced the two-fold axis from the DAMMIN model. A threading model of HC2 was generated from the structure of HC1 using Phyre (Kelley et al., 2015) and modelled along with bikunin (Xu et al., 1998) and HC1 into the DAMMIN envelope using Sculptor (Wahle and Wriggers, 2015) simultaneous docking protocols.

Structures of the rHC1 vWFa domain (this study) and the complement C3 C-terminal C345C domain (PDB 2XWJ (Forneris et al., 2010)) were positioned relative to each other informed by the C3-FB complex (2XWJ). The models were locally docked to each other using Rosetta_3.2 and the standard docking protocol (Leaver-Fay et al., 2011), with random perturbations of 3Å and 8°; 10,000 models were generated and the lowest energy model is shown in Figure 5D. Additionally, a SAXS envelope was generated of full length C3 bound to rHC1 in the presence of 2mM MnCl_2_. DAMMIN envelopes were calculated as described above. Crystal structures of C3 (PDB 2A73 (Janssen et al., 2005)) and HC1 (this study) were docked into this envelope using Sculptor (Wahle and Wriggers, 2015).

### Analytical ultracentrifugation of rHC1

The metal ion dependence of rHC1 dimerisation was analysed using both velocity and equilibrium AUC. All AUC experiments were conducted at 20°C on a Beckman XL-A ultracentrifuge with an An60Ti rotor.

For velocity AUC, 18µM WT rHC1 protein was prepared in HEPES buffered saline pH 7.5, either in the presence of 2.5mM EDTA or 5mM MgCl_2_. The samples were analysed at 40,000 rpm for 5h, with scans taken at 280nm every 90s. This experiment was conducted in triplicate with representative data shown in Figure 3A. Sedimentation coefficient distributions (c(s)) were calculated using SEDFIT (Schuck, 2000).

For equilibrium AUC, measurements were made at 3 different concentrations of rHC1 (4, 11, and 22 µM), where these were each prepared with 5 different concentration of MgCl_2_ (0 (2.5 mM EDTA), 0.1, 0.5, 1 and 5 mM). Rotor speeds of 10,000, 15,000, and 20,000 rpm were used with scans at 280nm (and 290nm for the highest concentration) after equilibrium had been reached (18 h). Data (from triplicate experiments) were analysed by global analysis with SEDFIT/SEDFAT (Houtman et al., 2007) and fitted to a monomer-dimer model.

### Surface plasmon resonance of rHC1-ligand interactions

SPR experiments were conducted on either a Biacore 3000 or T200 instrument. For C3 binding assays, ∼2,000 RU of C3 was immobilised on a CM5 chip using standard amide coupling chemistry and rHC1 was injected at a range of concentrations (1µM – 31.25µM) over the chip surface. For all other assays, ∼1,500 RU of rHC1 proteins (WT, D298A or ΔvWFa) were immobilised on a C1 chip by amide cross-linking chemistry and LAP (from TGFβ1; 3.125nM – 200nM), LTBP1 (NT1, EGF or CT domains; all at 0.156nM – 10nM), TGFβ1, TGFβ3 (both at 7.8nM – 500nM), TGFβ1-LAP, TGFβ2-LAP, TGFβ3-LAP (all at 3.125nM – 200nM) or vitronectin (0.3125nM – 10nM) were used as the analyte. Experiments were conducted in HEPES buffered saline, pH 7.5 with 0.05% (v/v) Tween-20. Metal ions (2mM) or chelating agent (EDTA; 10mM) were added to the buffers and a flow rate of 50 μl/min was used when generating kinetic parameters. Data were collected in triplicate and *K*_D_ values (mean ± S.D. in Table 3) were determined from multicycle kinetics, where data were fitted to a Langmuir 1:1 model using the BIAeval T200 software. For all fits, the Chi^2^ value obtained was less than 10% of the R_max_ value.

### C3 convertase assay

Inhibition of C3 activation to C3b was measured using a fluid phase convertase assay. Here C3 (19.5µM) was incubated with 1.75µM complement factor B (FB) and 0.37µM complement factor D in 20mM HEPES, 130mM NaCl, 3mM MgCl_2_, 1mM EGTA, pH 7.5. The effect of rHC1 (preincubated with 1mM MnCl_2_) was measured at concentrations ranging from 0 to 27µM; complement factor H (FH) was used as a positive control. After 1-min incubation at 37°C, the reaction was stopped by addition of 5x SDS loading buffer and samples were incubated at 100°C for 5 min. The samples were run on a 4-12% gradient SDS-PAGE gel and stained with Coomassie Blue. C3a formation was monitored by densitometry using an Odyssey imaging system (LI-COR Biosciences).

## Author Contributions (CRediT Compliant)

Conceptualization D.C.B. and A.J.D.; Methodology D.C.B., A.W.L-S. and T.A.J.; Formal analysis D.C.B.; Investigation D.C.B. and A.W.L-S.; Resources A.J.D., C.B., D.C.B., J.J.E., C.M.K., C.M.M.; Data curation D.C.B.; Writing – Original Draft D.C.B. and A.J.D.; Writing – Review & Editing all authors; Visualisation D.C.B.; Supervision A.J.D. and C.M.M.; Funding acquisition A.J.D. and C.M.M.

## Acknowledgements

We gratefully acknowledge funding from Arthritis Research UK (19489) and the Medical Research Council (K004441). We would also like to thank Ruth Steer and Helen Troilo for making the LTBP1 domains and the beamline scientists at Diamond (UK) and DESY PETRA III (Germany) where we collected crystallography (beamline IO3) and SAXS data (beamlines I22 and EMBL-P12). Other biophysical analyses were carried out in the BioMolecular Analysis Core Facility at the University of Manchester, which is supported by Centre funding from the Wellcome Trust (088785/Z/09/Z and 203128/Z/16/Z).

## Supplementary Figure Legends

**Supplementary Figure S1: Schematics of IαI and PαI, TSG-6-mediated HC transfer, and domain organisations for HCs 1-3. A)** A schematic illustrating the organisation of IαI and PαI showing that these proteoglycan both contain the bikunin core protein to which a chondroitin sulphate (CS) chain is attached via a typical tetrasaccharide linkage; heavy chains (HC1 and HC2 in IαI and HC3 in PαI) are linked to CS via ester bonds (red circles) formed between their C-terminal aspartic acid residues and a C6 hydroxyl within a N-acetyl galactosamine sugar in CS. In the presence of hyaluronan (HA), which is composed of a variable number (n) of repeating disaccharides of glucuronic acid (diamonds) and N-acetyl glucosamine (squares), and the trans-esterase TSG-6, HCs are covalently transferred from IαI/PαI onto HA to form HC•HA complexes; ester bonds link the C-terminal aspartic acids of the HCs to N-acetyl glucosamine residues of HA. **B)** A schematic of the domain organisation of the mature HC1 protein (residues 35-672 in Uniprot P19827), as determined from the crystal structure described here, along with their corresponding structural elements. The N- and C-terminal regions associate to form the Hybrid2 domain and the vWFA domain is flanked by H-sequences that constitute the Hybrid1 domain; the position of the D298A mutant is indicated along with the region that is deleted in the ΔvWFA construct. The domain organisations of the mature HC2 and HC3 proteins can be inferred from their homology with HC1 (39% and 54% identity, respectively).

**Supplementary Figure 2: Equilibrium AUC on rHC1 in absence and presence of MgCl_2_ ions.** Three concentrations of rHC1 (4, 11 and 22 μM) were analysed by equilibrium AUC at rotor speeds of 10,000 (pink), 15,000 (blue) and 20,000 (cyan) rpm in the absence (2.5 mM EDTA) or presence of 0.1, 0.5, 1 or 5mM MgCl_2_. High speed data for 11 μM HC1 in 1 mM MgCl_2_ were omitted.

**Supplementary Figure S3: SAXS data analysis for HC1 monomer (blue) and dimer (orange). A)** Analysis of the Guinier region and residuals. **B)** Dimensionless Kratky plots show that HC1 monomer and dimer molecules are folded and globular; the cross-hairs denote the globularity point, and the shift of the maxima for the HC1 dimer to the right indicate that it is more extended and asymmetric than the monomer. SIBYLS (**C**) and Porod-Debye (**D**) plots indicate that the HC1 monomer and dimer are rigid.

**Supplementary Figure S**4: **SAXS data analysis for IαI. A)** Analysis of the Guinier region and residuals. **B)** Dimensionless Kratky plots show that all of IαI is folded, globular, but asymmetric as revealed by the maxima being to the right of the globularity point (cross-hairs). SIBYLS (**C**) and Porod-Debye (**D**) plots demonstrate that IαI is rigid.

**Supplementary Figure S5**: **SAXS data analysis for the rHC1-C3 complex. A)** Analysis of the Guinier region and residuals. **B)** A Dimensionless Kratky plot reveals that the rHC1-C3 complex is folded, globular, but asymmetric as revealed by the maxima being to the right of the globularity point (cross-hairs). SIBYLS (**C**) and Porod-Debye (**D**) plots indicate that the rHC1-C3 complex has some flexibility.

**Supplementary Figure S6: MIDAS-independent binding of HC1 to vitronectin and TGFβ-LAP proteins.** SPR sensorgrams for the interaction of vitronectin (Vn), at concentrations of 10nM, 5nM, 2.5nM, 1.25nM, 0.625nM, with immobilised WT **(A)**, D298A **(B)** or ΔvWFa **(C)** rHC1. Data are representative of 3 independent experiments with derived numerical values shown in Table 3. **D)** SPR sensorgrams for the binding of TGFβ-LAP proteins (TGFβ1-LAP (green); TGFβ2-LAP (orange or red); TGFβ3-LAP (purple or blue)) with immobilised rHC1 (WT (light green, orange, purple) or D298A (dark green, red, blue)). The individual interactions were analysed further in 3 independent experiments (using different concentrations of TGFβ-LAP proteins) to generate the data in Table 3.

