## Supplementary figures and images for "Heavy chain-1 of inter-α-inhibitor has an integrin-like structure with immune regulatory activities"

### Figure S1

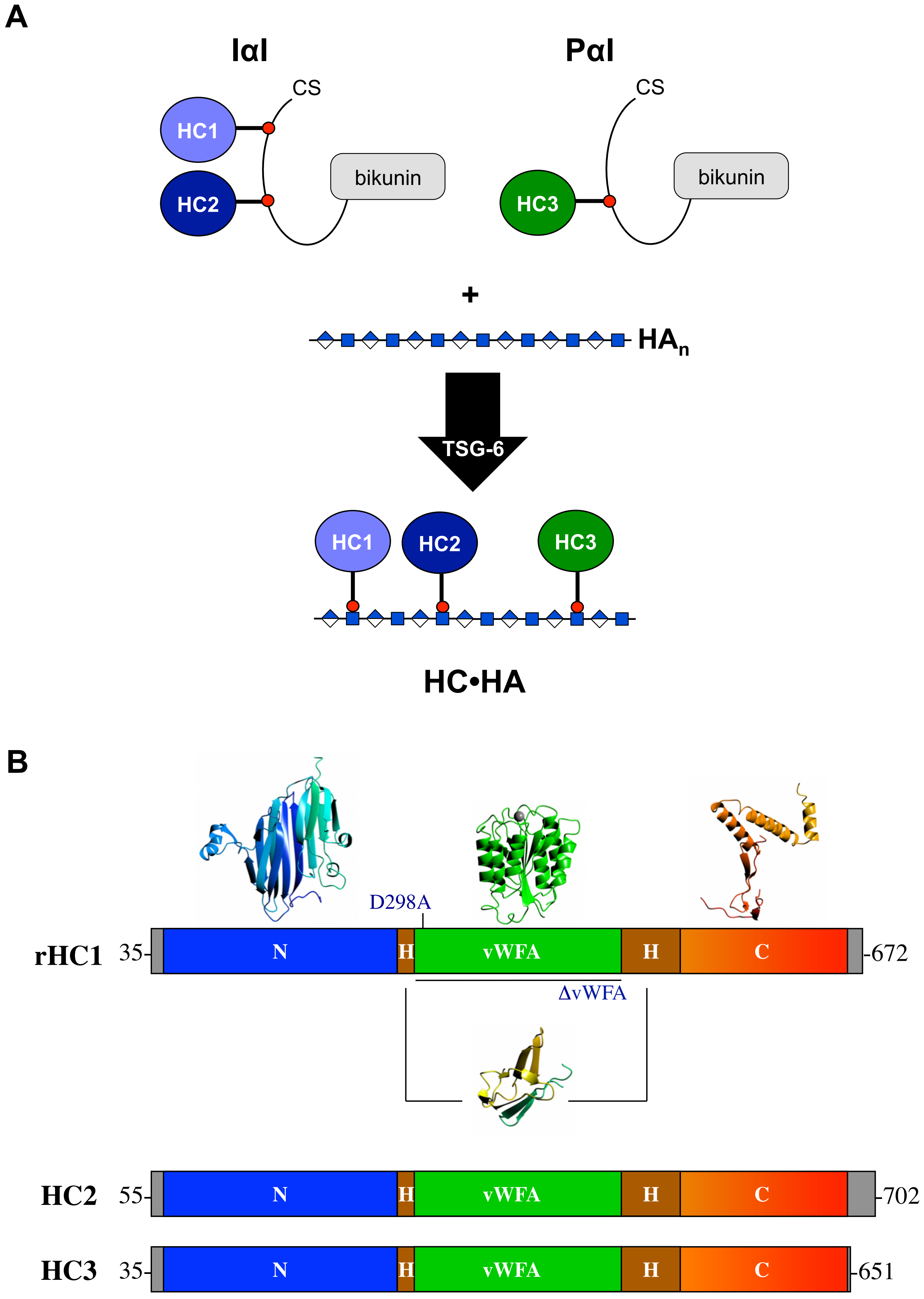

### Figure S2

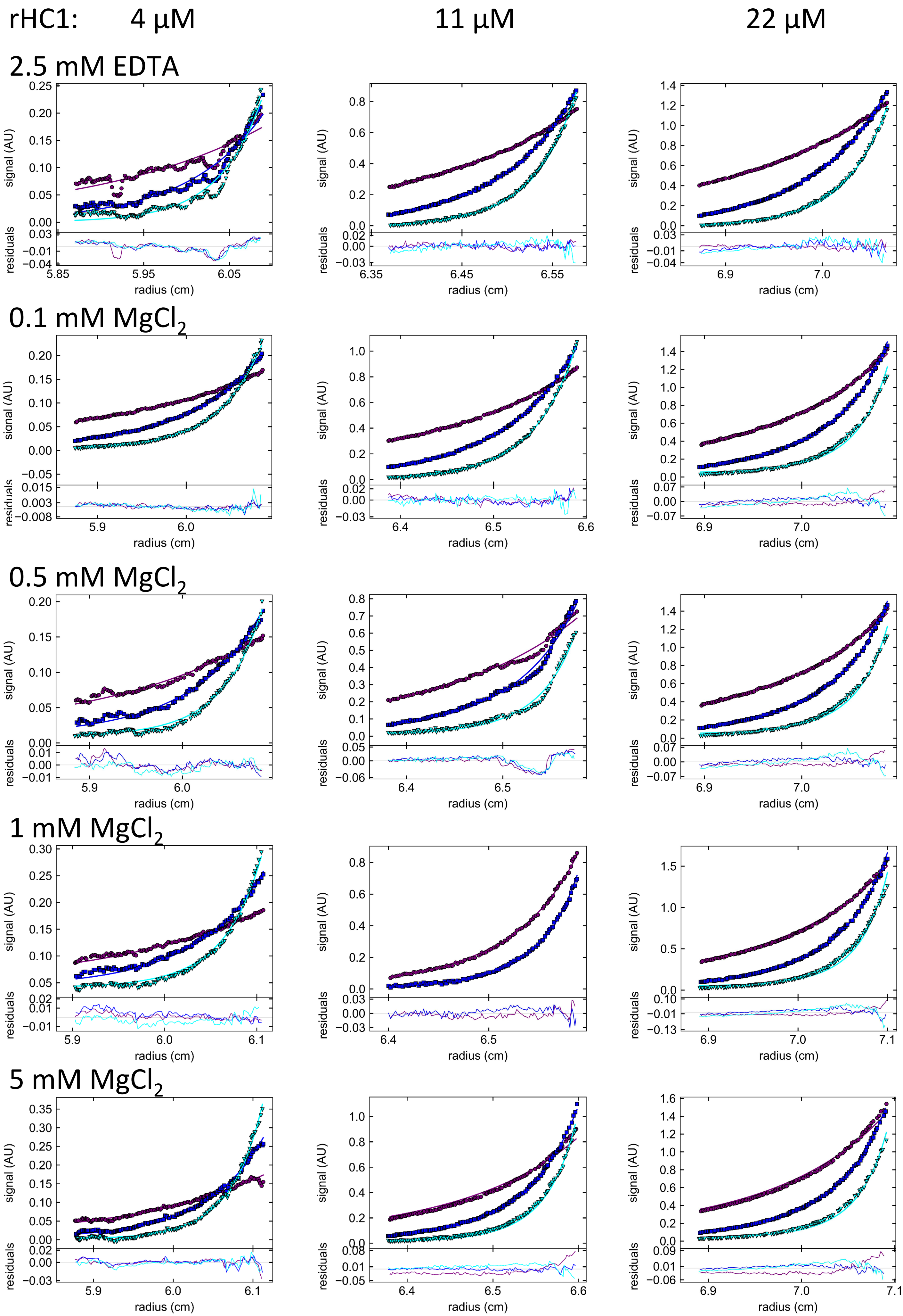

### Figure S3

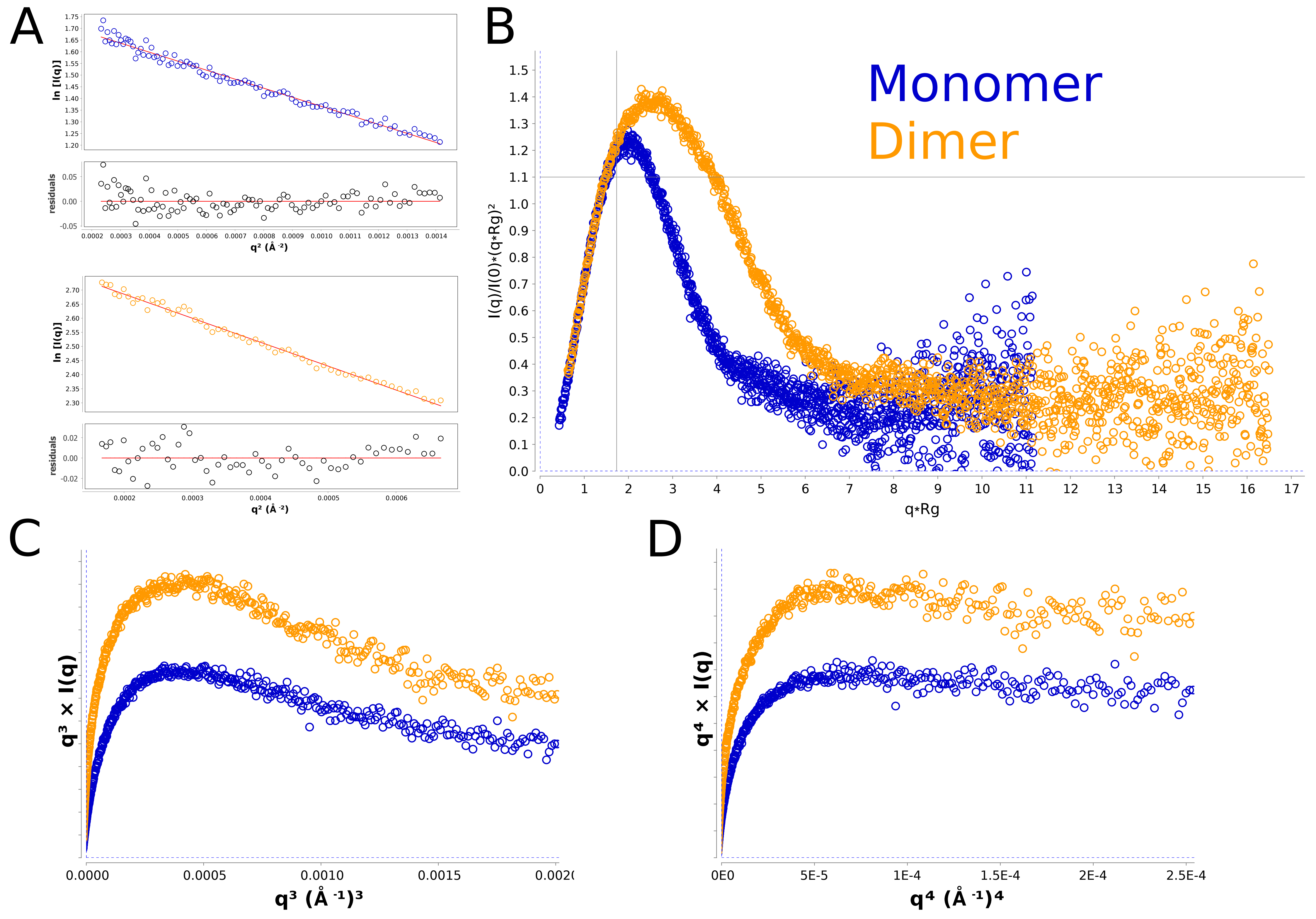

### Figure S4

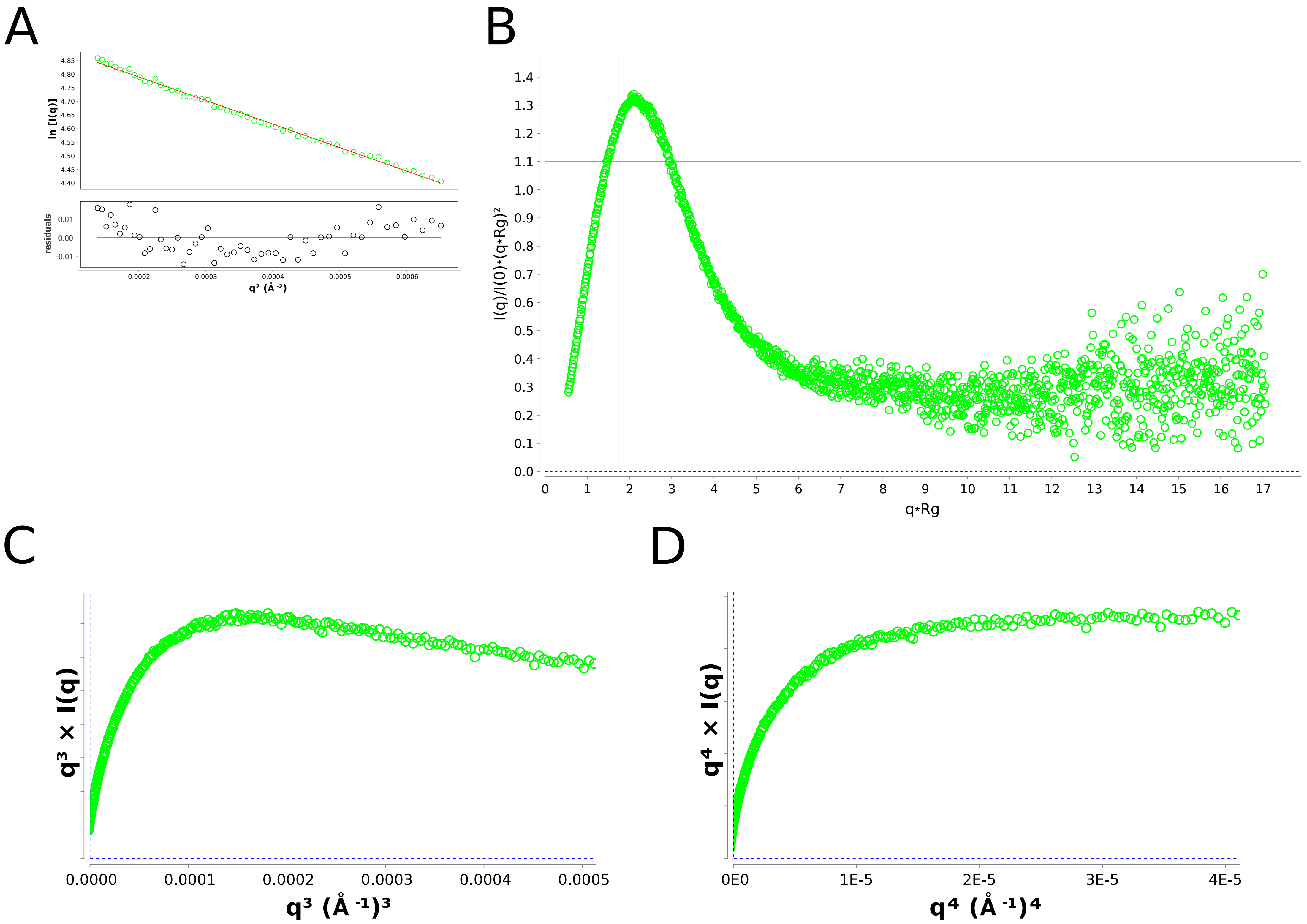

### Figure S5

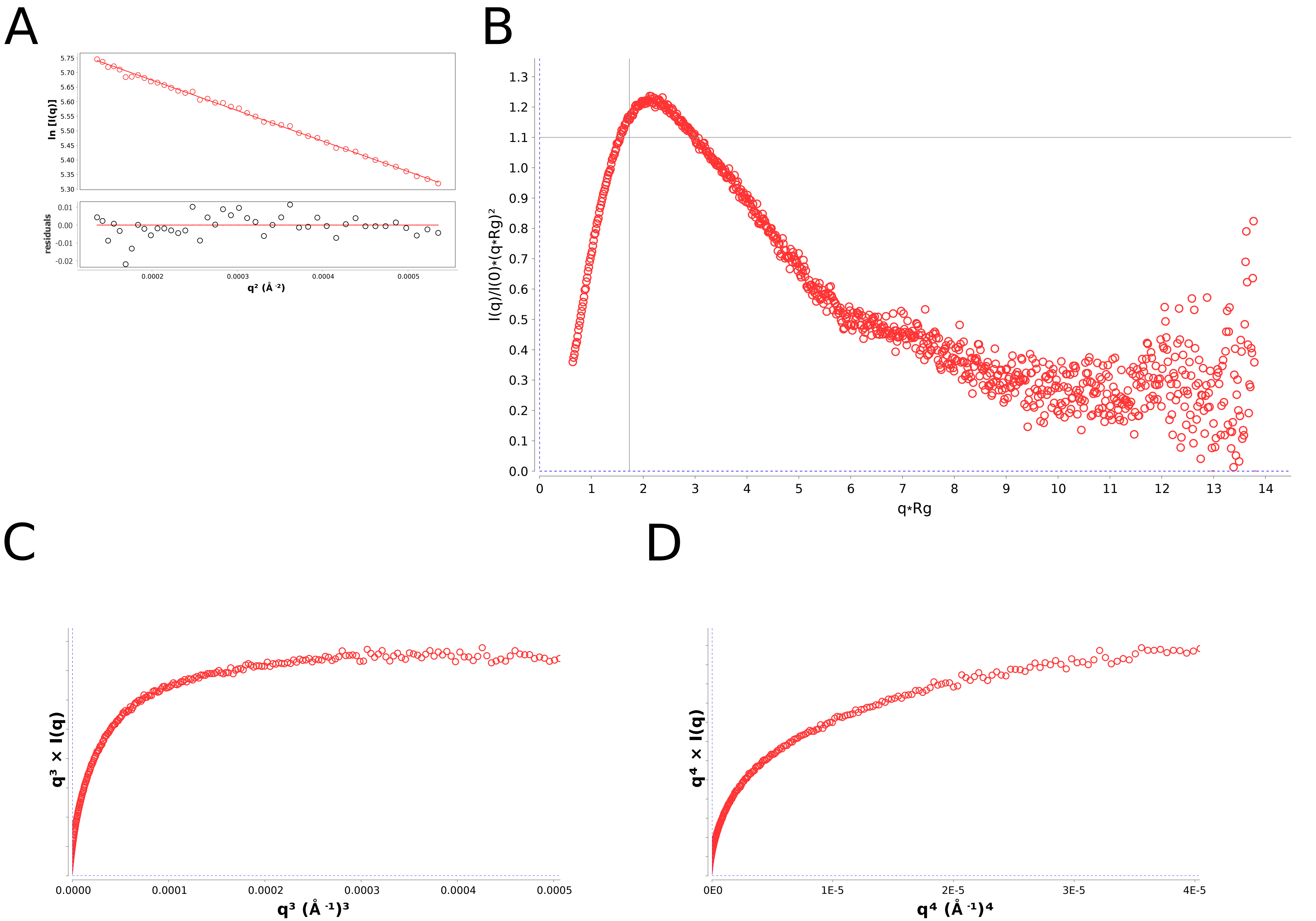

### Figure S6

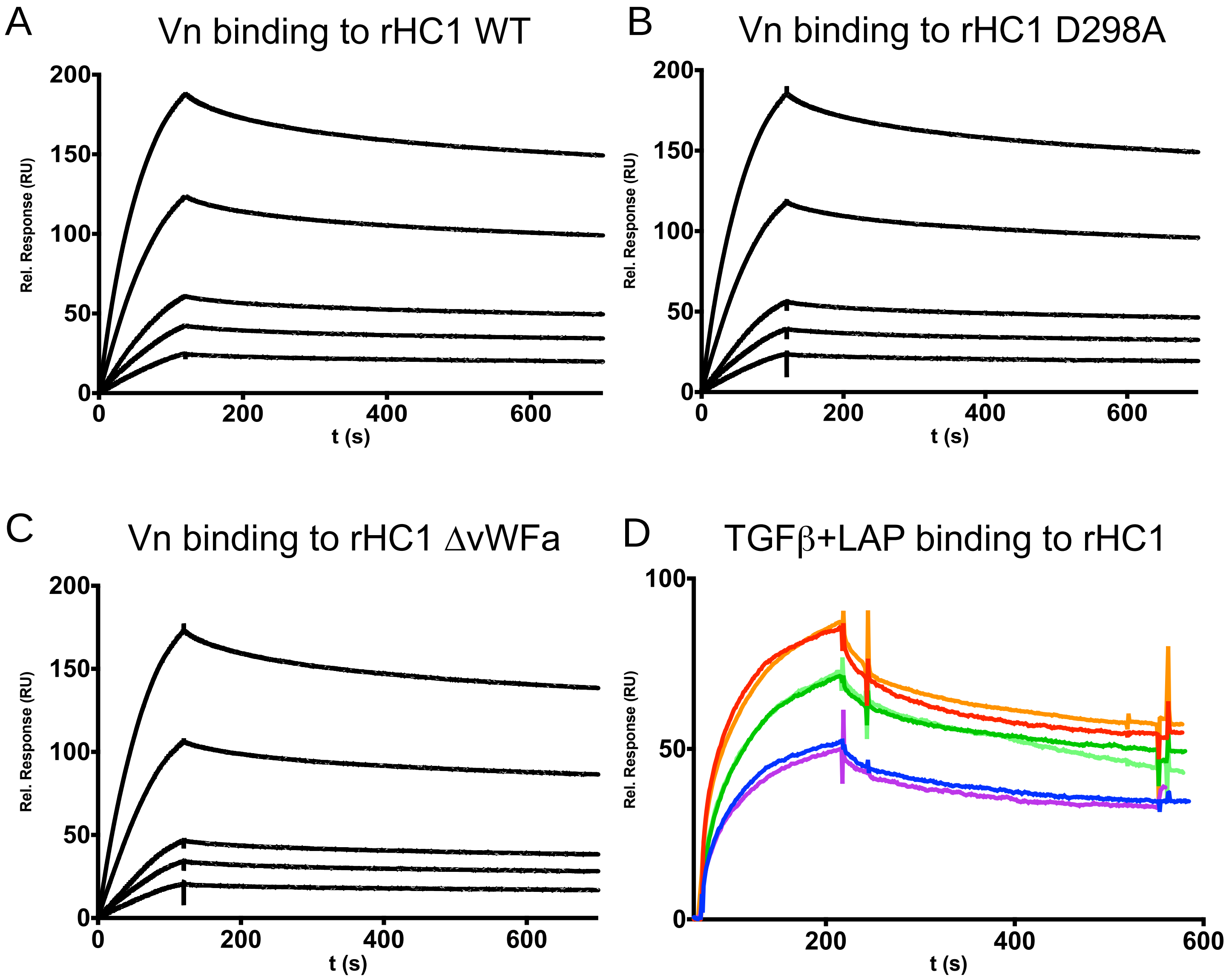
